# Developmentally regulated *Shh* expression is robust to TAD perturbations

**DOI:** 10.1101/609941

**Authors:** Iain Williamson, Lauren Kane, Paul S. Devenney, Eve Anderson, Fiona Kilanowski, Robert E. Hill, Wendy A. Bickmore, Laura A. Lettice

## Abstract

Topologically Associating Domains (TADs) have been proposed to both guide and constrain enhancer activity. *Shh* is located within a TAD known to contain all its enhancers. To investigate the importance of chromatin conformation and TAD integrity on developmental gene regulation, we have manipulated the *Shh* TAD – creating internal deletions, deleting CTCF sites including those at TAD boundaries, as well as larger deletions and inversions of TAD boundaries. Chromosome conformation capture and fluorescence in situ hybridisation assays were used the investigate changes in chromatin conformation that result from these manipulations. Our data suggest that the substantial alteration of TAD structure has no readily detectable effect on *Shh* expression patterns during development – except where enhancers are deleted - and results in no detectable phenotypes. Only in the case of a larger deletion of one TAD boundary could some ectopic influence of the *Shh* limb enhancer be detected on a gene - *Mnx1* in the neighbouring TAD. Our data suggests that, contrary to expectations, the developmental regulation of *Shh* expression is remarkably robust to TAD perturbations.

## Introduction

At the megabase-scale, the mammalian genome is partitioned into self-interacting topologically associated domains (TADs) (Dixon et al., 2012; Nora et al., 2012) Mammalian TAD boundaries are enriched in CTCF sites in a convergent orientation (Narendra et al., 2015; Rao et al., 2014; Sanborn et al., 2015). TADs are formed by dynamic cohesin-driven loop extrusion (Fudenberg et al., 2016; Nora et al., 2017; Rao et al., 2017; Schwarzer et al., 2017; Vian et al., 2018) and convergent CTCF sites act to impede loop extrusion allowing WAPL to release cohesin from the chromosome (Haarhuis et al., 2017).

The regulatory landscapes of developmental genes are frequently found to be contained together in the same TAD (Dixon et al., 2012; Rao et al., 2014). TADs have therefore been proposed to act as functional regulatory units that favour contacts between enhancers and their target gene within a TAD whilst limiting aberrant interactions of enhancers across TAD boundaries (Fudenberg et al., 2016; Sun et al., 2019). In support of this, some studies have found that deletion or inversion of CTCF sites at TAD boundaries, can promote TAD boundary crosstalk and re-wire enhancer-promoter contacts (de Wit et al., 2015; Guo et al., 2015; Narendra et al., 2015; Rodríguez-Carballo et al., 2017). Moreover, a number of recent studies have suggested that changes to TAD structure can disrupt gene regulation through enhancer-rewiring in human disease (Flavahan et al., 2016; Franke et al., 2016; Lupiáñez et al., 2015).

However, other studies report that, although depletion of CTCF erases the insulation between TADs, it has limited effects on gene expression (Nora et al., 2017; Soshnikova et al., 2010). To further study the CTCF mediated function of TADs in developmental gene regulation, we have exploited the sonic hedgehog (*Shh*) regulatory domain – a paradigm locus for long-range regulation. The SHH morphogen controls the growth and patterning of many tissues during embryonic development, including the brain, neural tube and the limb. Spatial and temporal *Shh* expression is regulated by tissue-specific enhancers located within the gene, and in a large gene desert upstream of the gene (Anderson and Hill, 2014). *Shh* and its *cis*-acting elements are all contained within a well-characterised ∼960kb TAD (Anderson et al., 2014; Williamson et al., 2016). In the developing limb bud, *Shh* expression is solely determined by the ZRS enhancer (Sagai, 2005) located within an intron of the widely expressed *Lmbr1*, located 850kb upstream of *Shh*. Fluorescence *in situ* hybridisation (FISH) has shown that *Shh* and the ZRS are consistently located in relatively close proximity to each other in all cell types and tissues examined which we infer to be a consequence of the underlying invariant TAD structure. In contrast, we have observed increased ZRS-*Shh* colocalisation only in the *Shh*-expressing posterior portion of developing limb buds (Williamson et al., 2016). This might be consistent with a specific gene-enhancer contact.

To investigate the importance of chromatin architecture on TAD structure and thus on the regulation of gene expression, here, we extensively manipulate the *Shh* TAD and its TAD boundaries. We use a chromosome conformation assay (5C) and FISH to investigate how these manipulations affect TAD structures and interactions within and between TADs and we determine how these alterations affect the expression pattern of *Shh* and other developmentally regulated genes nearby. We also examine the phenotypic consequences of these manipulations. Our results question how important TADs are for correct spatial and temporal gene regulation.

## Materials and Methods

### Cell Culture and CrispR/cas9 mediated deletions

E14TG2A mouse embryonic stem cells (ESCs) were cultured under standard conditions (Anderson et al., 2014). CrispR guides were made by cloning annealed oligos (Table S1) into px458 (Addgene). 2μg of vector DNA were transfected into 8×10^5^ ESCs using Lipofectamine 2000 (ThermoFisher) following the manufacturer’s instructions. After 48 hours, GFP positive cells were sorted by FACS and plated at low density. Ten days later, individual clones were picked and screened for correct deletion by PCR and Sanger sequencing (primers are listed in supplementary material Table S1).

### Mouse lines and embryo analysis

The *Shh*^Δ700^ deletion was created by crossing the line SBLac96 (Anderson et al., 2014) to a line carrying a pCAGGS-Cre recombinase gene (Araki et al., 2006). With the exception of the Δ35kb and Inv35kb mouse lines, which were made by injection of the ESCs in to blastocysts, all of the other mouse lines were created as in Lettice et al., (2017) by direct microinjection into C57Bl6/ CBA F2 zygotes of the same guides as were used in ESCs. Resultant G0 mice are screened by PCR using flanking primers (Supp Table 1) and the deletions confirmed by Sanger sequencing. Lines were then established by crossing founder mice to C57Bl6 wildtypes.

LacZ expression analysis, in situ hybridisations and RT-PCR reactions were conducted as in Anderson et al. (2014).

### FISH

E11.5 embryos were collected, fixed, embedded, sectioned, antibody stained for SHH expression and processed for FISH as previously described (Morey et al., 2007, Lettice et al., 2014), except that sections were cut at 8 μm. Fosmid clones (Figure 1A, Table S3) were prepared and labelled as previously described (Morey et al., 2007). Between 160-240 ng of biotin- and digoxigenin-labelled fosmid probes were used per slide, with 16-24 μg of mouse Cot1 DNA (Invitrogen) and 10 μg salmon sperm DNA. For 4-colour FISH, similar quantities of the additional fosmid was labelled with either Green496-dUTP (Enzo Life Sciences) or red-dUTP (Alexa Fluor^TM^ 594-5-dUTP, Invitrogen).

**Figure 1.**
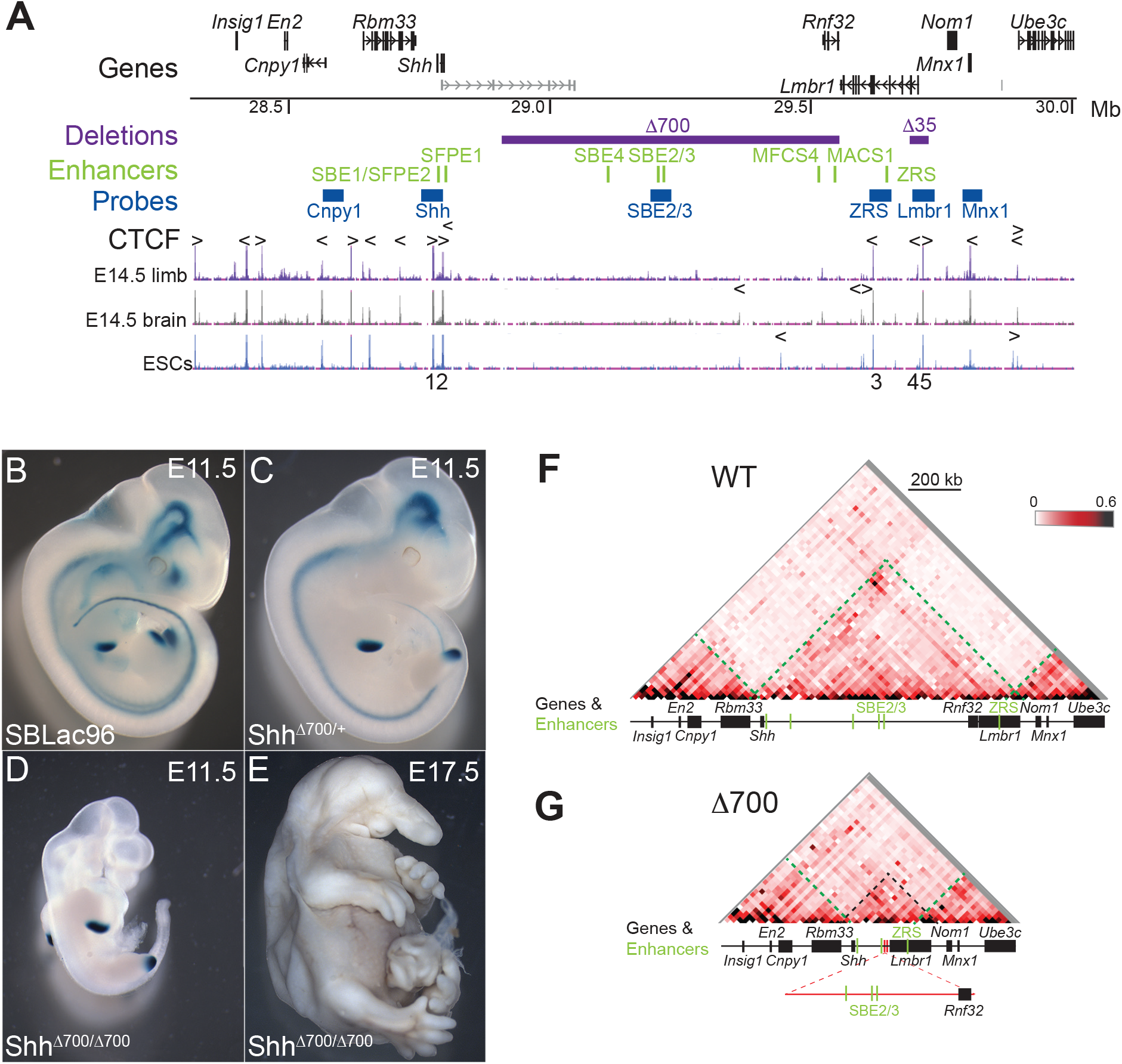
A 700-kb intra-TAD deletion has no adverse effects on limb-specific expression of *Shh*. (**A**) (Top) Location of genes over a 1.7 Mb murine genomic locus (chr5: 28317087-30005000; mm9) containing *Shh* analysed by 5C, with the position of tissue-specific *Shh* enhancers shown below in green. Locations to which the fosmid probes used for FISH hybridize are shown in blue, and the purple bars indicate deleted genomic regions (Δ700 and Δ35). The bottom three tracks show UCSC ENCODE CTCF ChIP-seq tracks displaying the CTCF binding profile in E14.5 limb buds and brain and in ESCs. Arrowheads above the tracks indicate the orientation of CTCF-binding motifs and the deleted CTCF binding sites are numbered below. **(B-D)** Staining for the LacZ gene carried by the sleeping beauty transposon in E11.5 embryos, **(B)** carries the intact SBLac96 chromosome while **(C)** shows the remaining sites of expression after cre mediated deletion of 700Kb and **(D)** shows the phenotype of an embryo homozygous for the 700kb deletion, with LacZ staining only evident in the ZPA of the limb buds. A homozygous deletion embryo **(E)** at E17.5 showing craniofacial and brain defects but normally formed limbs. Heat-maps showing 5C data from cells of the bodies of E11.5 wild-type embryos **(F)** and embryos homozygous for the 700kb deletion **(G)**, across the 1.7-Mb *Shh* region shown in **(A)**. Heat map intensities represent the average of interaction frequency for each window, colour-coded according to the scale shown. Interaction frequencies were normalized based on the total number of sequence reads in the 5C data set and the data shown is binned over 22.5-kb windows. Green dashed lines highlight the TAD boundary locations, black dashed lines indicate the *Shh* TAD boundaries and the reduced size of the TAD. Data for biological replicates are in Supplemental Figure S1.

For 3D FISH on ESCs, 1×10^6^ cells were seeded on slides for overnight. Cells were fixed in 4% paraformaldehyde (pFA) for 10 mins at room temperature and then permeabilized using 0.5% TritonX for 10 mins (Eskeland et al., 2010).

### Image analysis

Slides were imaged using a Photometrics Coolsnap HQ2 CCD camera and a Zeiss AxioImager A1 fluorescence microscope with a Plan Apochromat 100x 1.4NA objective, a Nikon Intensilight Mercury based light source (Nikon UK Ltd, Kingston-on-Thames, UK) and either Chroma #89014ET (3 colour) or #89000ET (4 colour) single excitation and emission filters (Chroma Technology Corp., Rockingham, VT) with the excitation and emission filters installed in Prior motorised filter wheels. A piezoelectrically driven objective mount (PIFOC model P-721, Physik Instrumente GmbH & Co, Karlsruhe) was used to control movement in the z dimension. Step size for z stacks was set at 0.2 μm. Hardware control, image capture and analysis were performed using Nikon Nis-Elements software (Nikon UK Ltd, Kingston-on-Thames, UK). Images were deconvolved using a calculated point spread function with the constrained iterative algorithm of Volocity (Perkinelmer Inc, Waltham, MA). Image analysis was carried out using the Quantitation module of Volocity (Perkinelmer Inc, Waltham, MA).

### 3C library preparation

Limbs buds and bodies (with the limbs and heads removed) from wild type embryos, and entire *Shh*^Δ700/Δ700^ embryos were dissected at E11.5 and the tissue dissociated by pipetting in just enough PBS to cover them. The cells were fixed with 1% formaldehyde for 10 min at room temperature. For ESCs, 5×10^6^ – 1×10^7^ cells were fixed. Crosslinking was stopped with 125 mM glycine, for 5 min at r.t. followed by 15 min on ice. Cells were centrifuged at 400 *g* for 10 min at 4°C, supernatants removed, and cell pellets flash frozen on dry ice before storage at −80°C.

Cell pellets were treated as previously described (Dostie and Dekker, 2007; Ferraiuolo et al., 2010; Williamson et al., 2014). HindIII-HF (NEB) was the restriction enzyme used to digest the crosslinked DNA.

### 5C primer and library design

5C primers covering the *Usp22* (mm9, chr11: 60,917,307-61,003,268) and *Shh* regions (mm9, chr5: 28,317,087-30,005,000) were designed using ‘my5C.primer’ (Lajoie et al., 2009) with the following parameters: optimal primer length of 30 nt, optimal TM of 65°C, default primer quality parameters (mer:800, U-blast:3, S-blasr:50). Primers were not designed for large (>20 kb) and small (<100 bp) restriction fragments, for low complexity and repetitive sequences, or where there were sequence matches to >1 genomic target. The *Usp22* region was used to assess the success of each 5C experiment but was not used for further data normalization or quantification.

The universal A-key (CCATCTCATCCCTGCGTGTCTCCGACTCAG-(5C-specific)) and the P1-key tails ((5C-specific)-ATCACCGACTGCCCATAGAGAGG) were added to the Forward and Reverse 5C primers, respectively. Reverse 5C primers were phosphorylated at their 5′ ends. An alternating design consisting of 365 primers in the *Shh* region (182 Forward and 183 Reverse primers) was used. Primer sequences are listed in Table S9.

### 5C library preparation

5C libraries were prepared and amplified with the A-key and P1-key primers as described in (Fraser et al., 2012). Briefly, 3C libraries were first titrated by PCR for quality control (single band, absence of primer dimers, etc.), and to verify that contacts were amplified at frequencies similar to that usually obtained from comparable libraries (same DNA amount from the same species and karyotype) (Dostie and Dekker, 2007; Dostie et al., 2007; Fraser et al., 2010). We used 1 - 10 μg of 3C library per 5C ligation reaction.

5C primer stocks (20 μM) were diluted individually in water on ice and mixed to a final concentration of 2 nM. Mixed diluted primers (1.7 μl) were combined with 1 μl of annealing buffer (10X NEBuffer 4, New England Biolabs Inc.) on ice in reaction tubes. 1.5 μg salmon testis DNA was added to each tube, followed by the 3C libraries and water to a final volume of 10 μl. Samples were denatured at 95°C for 5 min and annealed at 55°C (48°C ESCs) for 16 hours. Ligation with Taq DNA ligase (10 U) was performed at 55°C (48°C ESCs) for one hour. One tenth (3 μl) of each ligation was then PCR-amplified individually with primers against the A-key and P1-key primer tails. We used 26 cycles based on dilution series showing linear PCR amplification within that cycle range. The products from 3 to 5 PCR reactions were pooled before purifying the DNA on MinElute columns (Qiagen).

5C libraries were quantified by bioanalyser (Agilent) and diluted to 26 pmol (for Ion PGM™ Sequencing 200 Kit v2.0). One microlitre of diluted 5C library was used for sequencing with an Ion PGM™ Sequencer. Samples were sequenced onto Ion 316™ Chips following the Ion PGM™ Sequencing 200 Kit v2.0 protocols as recommended by the manufacturer (Life Technologies^TM^).

### 5C data analysis

Analysis of the 5C sequencing data was performed as described in (Berlivet et al., 2013). The sequencing data was processed through a Torrent 5C data transformation pipeline on Galaxy (https://main.g2.bx.psu.edu/). Before normalizing, interactions between adjacent fragments were removed due to the high noise: signal ratio likely to occur here. Data was normalized by dividing the number of reads of each 5C contact by the total number of reads from the corresponding sequence run. All scales shown correspond to this ratio multiplied by 10^3^. The number of total reads and of used reads is provided for each experiment in Table S10. 5C datasets are to be uploaded to the Gene Expression Omnibus (GEO) website http://www.ncbi.nlm.nih.gov/geo/.

## RESULTS

### A large deletion within the *Shh* TAD does not disrupt local genome organisation or limb-specific activation of *Shh*

Prominent architectural features of the *Shh* TAD include CTCF bound at five binding sites at both boundaries across multiple cell types (Fig. 1A), and two sub-TADs with overlapping boundaries located within the gene desert between the forebrain enhancers and *Rnf32* (Figs. 1F and 2A heat maps). This region of the gene desert includes less well defined CTCF peaks that are not invariant across cell types but due to their location may have some role in defining these sub-TADs (Fig. 1A) (Rosenbloom et al., 2013).

To determine the contribution of TAD internal sequence to 3D chromatin organisation and gene expression, we exploited our previous work that used the local hopping activity of the sleeping beauty (SB) transposon to probe the *Shh* regulatory domain (Anderson et al., 2014). Transposition of the SB leaves a LoxP site at the initial integration site and inserts a second LoxP site where it re-integrates, enabling Cre recombinase to create deletions of the intervening DNA. The orientation of the re-integration means the LacZ gene carried by the SB is retained in the deleted chromosome allowing remaining enhancer activity to be monitered. Using this approach, we deleted approximately 700kb (∼70%) of the internal *Shh* TAD sequence, including the sub-TAD boundaries, but, leaving the five CTCF binding sites at the extremes of the TAD intact (Fig. 1A). The Δ700 deletion removes many of the known *Shh* enhancers, and relocates the ZRS to within 96kb of the *Shh* promoter (Fig. 1A). Removal of the *Shh* forebrain and epithelial enhancers in the Δ700 deletion is shown by LacZ staining of *Shh*^Δ700/+^ embryos which shows staining only within the floor plate and hind brain, presumably driven by the proximal enhancers SFPE1/2 and SBE1, and within the limbs driven by the ZRS (Figs. 1B & C). Homozygous *Shh*^Δ700/Δ700^ embryos show phenotypes very similar to those of *Shh*^-/-^ embryos but with normal limb and digit patterning (Chiang et al., 1996). These data indicate that, despite its incorrect position now only 96kb from the *Shh* promoter, ZRS is able to function normally to drive *Shh* expression in limb development (Figs. 1D & 1E).

To determine how removal of this ∼700kb disrupts local TAD structure we carried out 5C on whole E11.5 *Shh* ^Δ700/Δ700^ and wild type embryos. 5C heatmaps show that the *Shh* TAD boundaries and the adjacent TADs are unaffected by the deletion, (Figs. 1F, 1G & S1). Together, these data show that interactions within the deleted region are not required for ZRS activity or for maintaining the location of the TAD boundaries, and there is not a requirement for a great genomic distance between *Shh* and its limb enhancer.

### Interactions within the *Shh* TAD are delineated by CTCF sites either side of *Shh* and within *Lmbr1*

Our previous 5C analyses on cells dissected from whole limbs, bodies and heads of E11.5 embryos showed enriched interactions between the genomic region containing *Shh*, located at one TAD boundary, and a genomic region within *Lmbr1* close to ZRS, located ∼70kb from the other TAD boundary (Williamson et al., 2016). That this enrichment can be identified throughout the E11.5 embryo, a stage when we have shown that high levels of *Shh*-ZRS colocalisation occur only in the posterior distal limb, excludes active *Shh*-ZRS colocalisation as the sole driver of this apparent chromatin loop (Williamson et al., 2016). To gain further insight into the nature of these interactions, we dissected E11.5 limb buds to compare cell populations with no ZRS activity (anterior 2/3 of bud) with those with ZRS-active cells (posterior 1/3) (Figs. 2A and S2A).

The *Shh* TAD structure revealed by 5C is similar in both anterior and posterior limb bud cell populations, and comparable to dissected E11.5 bodies (compare Fig. 1F and Fig. 2A). At high (15kb) resolution, the strongest enrichment in both populations involved interactions between both *Shh* and the genomic region immediately 3′ of *Shh*, and a locus ∼20kb from ZRS in intron 5 of *Lmbr1* (Figs. 2B & S2B, left- and right-hand heatmaps). ENCODE data (Rosenbloom et al., 2013) indicates these three loci are all bound by CTCF across a range of cell and tissue types (Fig. 1A), with the underlying DNA containing CTCF-binding motifs in a convergent orientation suggestive of roles in blocking loop extrusion (Fig. 2A). There is a subtle enrichment of interactions apparently between *Shh* and ZRS in the posterior population which contains ZRS-active cells, known as the Zone of Polarising Activity (ZPA) (Figs. 2B & S2B, centre heatmap). The minimal difference between the anterior and posterior tissue could be due to the presence of cells in the posterior population where ZRS is not active diluting the interactions between *Shh* and ZRS.

**Figure 2.**
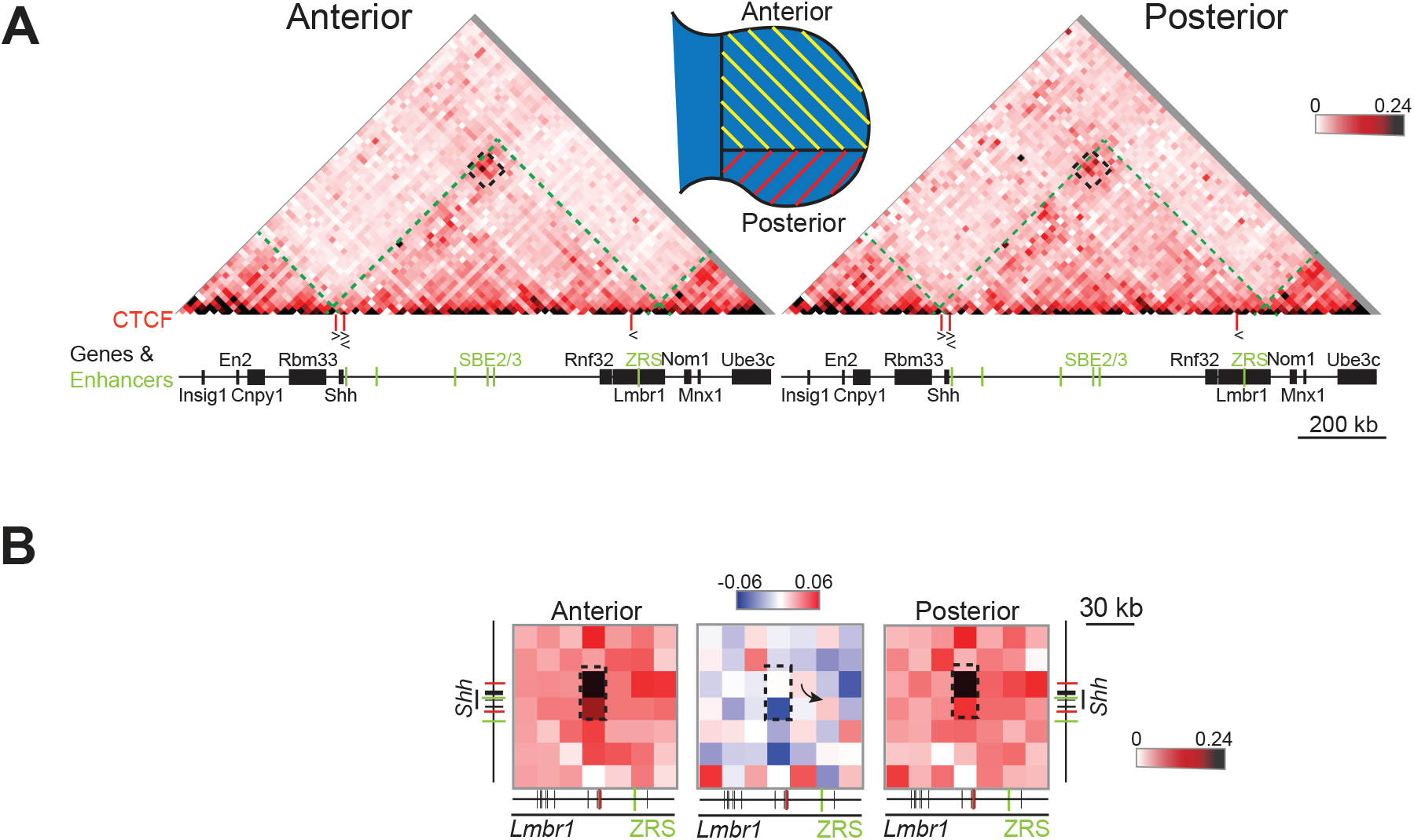
5C analysis in E11.5 distal anterior and posterior limb tissue. (**A**) 5C heat maps showing data from distal anterior and posterior limb bud cells of E11.5 embryos, across the 1.7-Mb *Shh* region shown in Figure 1A. Heat map intensities represent the average interaction frequency for each window, colour-coded according to the scale shown. Interaction frequencies were normalized based on the total number of sequence reads in the 5C data set and the data shown is binned over 22.5-kb windows. Green dashed lines indicate TAD boundaries, the interactions highlighted by the black dashed boxes locate the region of the heat maps shown in **(B)** at higher resolution. The schematic indicates the limb bud portions dissected for anterior and posterior cell populations. **(B)** Higher resolution (15kb binning) 5C heat maps from data displayed in **(A)** showing interactions between 105kb genomic regions encompassing *Shh* and ZRS. Left-hand and right-hand heat maps from anterior and posterior tissues respectively with intensities representing the average of interaction frequency for each window, colour-coded according to the scale shown. The comparison heatmap (centre) shows interactions enriched in posterior cells (red) or anterior cells (blue). Enriched interactions between loci containing CTCF binding sites are indicated by the black dashed boxes, the arrow in the comparison heat map highlights interactions between *Shh* and ZRS. Data for biological replicates are in Supplemental Figure S2.

### Deletion of CTCF sites at *Shh* reduces *Shh* intra-TAD interactions and disrupts *Shh*/ZRS proximity

The CTCF-anchored loop located at convergent binding sites near to both *Shh* and ZRS detected by 5C throughout the E11.5 embryo would appear to be important for maintaining *Shh*/ZRS spatial proximity and possibly contributes to the formation of the *Shh* TAD along with the CTCF binding sites located at the *Lmbr1* promoter (Fig. 1A). We therefore used CRISPR-Cas9 to delete sequences containing these domains of CTCF binding in mouse ESCs (The size for each individual deletion is listed in Table S1). We first deleted the CTCF binding regions 3′ and 5′ of *Shh* (sites numbered 1 and 2 respectively in Fig. 1A). We generated ESC lines homozygous for each of these deletions and compared chromatin conformation by 5C and FISH.

The *Shh* TAD structure in ESCs is similar to that in E11.5 embryos (Figs. 3A & S3A, top row, left-hand 5C heatmaps). Deletion of CTCF site 1 (ΔCTCF1), which delineates the TAD boundary 3′ of *Shh*, re-locates the TAD boundary by ∼40kb to 5′ of *Shh* to the vicinity of CTCF2 (Figs. 3A & S3A, top row, centre heatmaps). This is evidenced by *Shh* losing interactions with the rest of its own TAD in ΔCTCF1 cells, and gaining interactions with regions just 5′ of *En2* and *Rbm33* (Figs. 3A & S3A, bottom row, left-hand heatmaps). There is also loss of interactions with a locus upstream of the forebrain enhancers near the sub-TAD boundary within the larger *Shh* TAD. These data would be consistent with CTCF1 forming the *Shh* TAD boundary by blocking a loop extrusion process emanating from within the *En2* TAD.

**Figure 3.**
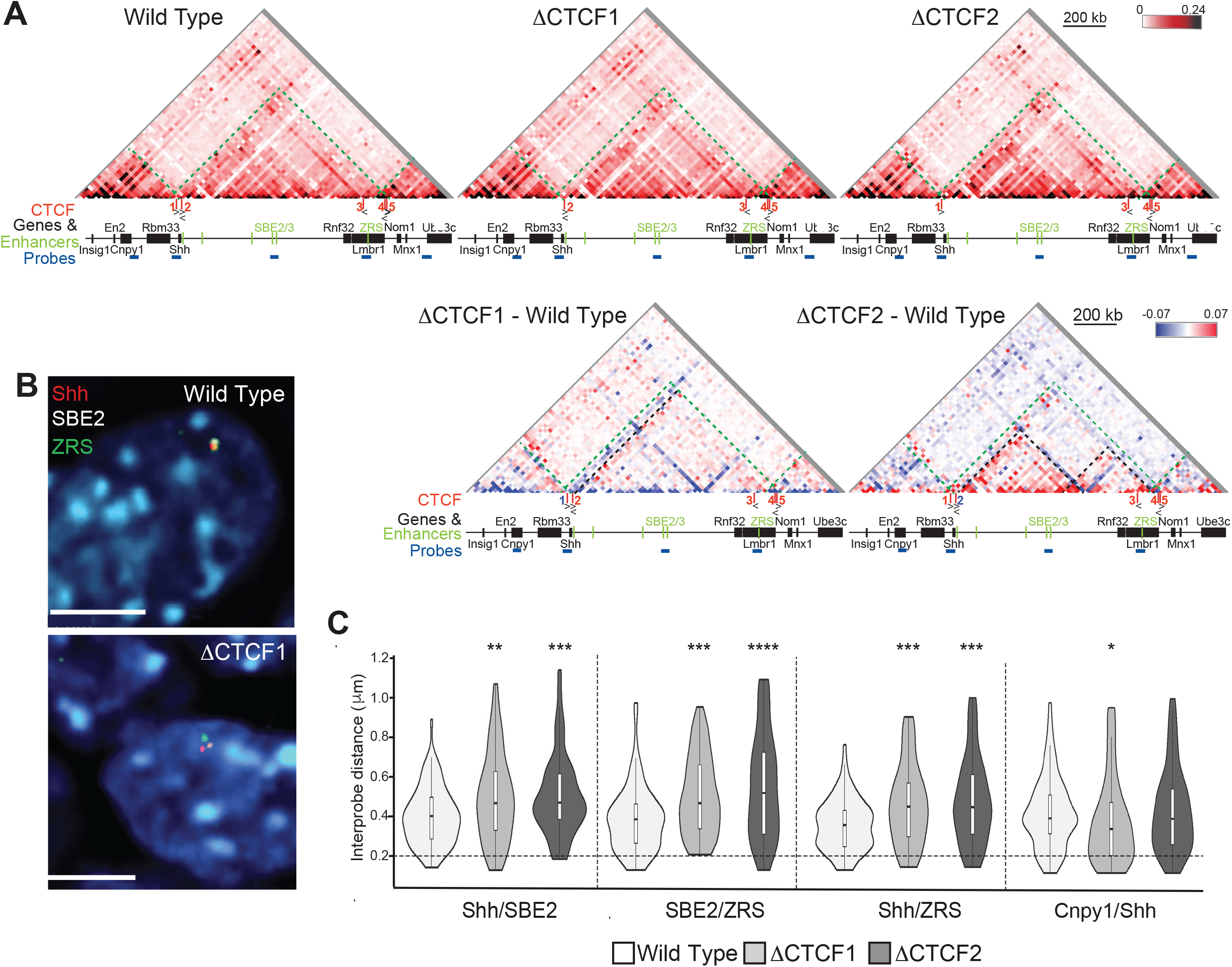
5C and 3D FISH identifies perturbations to chromatin conformation in ΔCTCF1 and ΔCTCF2 ESCs. **(A)** 5C heat maps showing data from wild type, ΔCTCF1 and ΔCTCF2 ESCs (top), across the 1.7-Mb *Shh* region shown in Figure 1A. Heat map intensities represent the average of interaction frequency for each window, colour-coded according to the scale shown. Interaction frequencies were normalized based on the total number of sequence reads and the data shown is binned over 22.5-kb windows. Below are heat maps comparing ΔCTCF1 or ΔCTCF2 enrichment (red) with wild type (blue). Green dashed lines indicate TAD boundaries, black dashed lines highlight change in TAD boundary position (ΔCTCF1) or enriched contacts within sub-TADs (ΔCTCF2). Data for biological replicates are in Supplemental Figure S3. **(B)** Images of representative nuclei from wild type and ΔCTCF1 ESCs showing FISH signals for *Shh*/SBE2/ZRS probes. Scale bars = 5 μm. **(C)** Violin plots show the distribution of interprobe distances (μm) between *Shh*/SBE2, SBE2/ZRS and *Shh*/ZRS probes in wild type, ΔCTCF1 and ΔCTCF2 ESCs. Horizontal dashed line shows the proportion of alleles that are colocalised (< 200 nm). The statistical significance between data sets was examined by Mann-Whitney U Tests, * < 0.05, ** < 0.01, *** < 0.001, **** < 0.0001.

The left hand *Shh* TAD boundary is not affected by deletion of CTCF site 2 (Figs. 3A & S3A, top row, right-hand heatmaps), however, the 5′ *Shh* region does gain contacts with the *En2* TAD in a similar manner to the loss of CTCF1 (Figs. 3A & S3A, bottom row, right-hand heatmaps), suggesting that both CTCF1 and CTCF2 are necessary to optimally block loop extrusion emanating from the *En2* TAD. There are also strongly enriched interactions within the *Shh* sub-TADs detected in ΔCTCF2 cells (Figs. 3A & S3A).

We also analysed possible alterations of chromosome conformation due to the CTCF site deletions with 3D-FISH using probes for *Shh*, SBE2 and ZRS (Fig. 3B). Interprobe distances between all three probe pairs were significantly increased in the CTCF deletion cells compared to wild type ESCs (Fig. 3C), consistent with the reduced interactions between *Shh* and the rest of the TAD identified by 5C. Conversely, we detected significantly decreased distances between *Shh* and *Cnpy1* (in the neighbouring *En2* TAD) in ΔCTCF1 cells (but not ΔCTCF2) compared to wild type (Figure 3C), consistent with the relocation of the TAD boundary.

Deleting either CTCF1 or CTCF2 disrupts *Shh*-ZRS spatial proximity in ESCs and, more generally, result in reduced 5C interactions between *Shh* and the rest of the regulatory TAD that may be due to the re-location of the TAD boundary (ΔCTCF1) or greater sub-division of the TAD (ΔCTCF2). The TAD boundary adjacent to *Shh* is sharply defined by CTCF1 whereas the boundary location of the neighbouring *En2* TAD cumulatively results from both CTCF 1 and 2, possibly by blocking loop extrusion emanating from this TAD. However, neither of these deletions on their own is sufficient to cause the merging of the two and neighbouring TADs.

### *Shh*-ZRS proximity is disrupted by the deletion of ZRS/*Lmbr1* CTCF sites

Both CTCF1 and CTCF2 have highly enriched interactions with the CTCF site ∼20 kb from ZRS in intron 5 of *Lmbr1* (CTCF3) (Figure 1A, site 3). Therefore, we deleted both alleles of CTCF3 (ΔCTCF3) to determine the consequences for chromosome conformation.

Whilst whole TAD integrity was unaffected by ΔCTCF3 (Figs. 4A & S4A, top row, centre-left heatmaps), intra-TAD reorganisation occurred in a similar manner to the loss of CTCF2, with enriched interactions within the sub-TADs (Figs. 4A & S4A). FISH inter-probe distances between *Shh*, SBE2 and ZRS (Fig. 4B) all significantly increased in ΔCTCF3 cells compared to wild type (Fig. 4C). These data suggest that loss of any one of the three CTCF binding domains (1, 2 or 3) can disrupt the spatial proximity of *Shh*, SBE2 and ZRS (Fig. 4C, Figure 3C).

**Figure 4.**
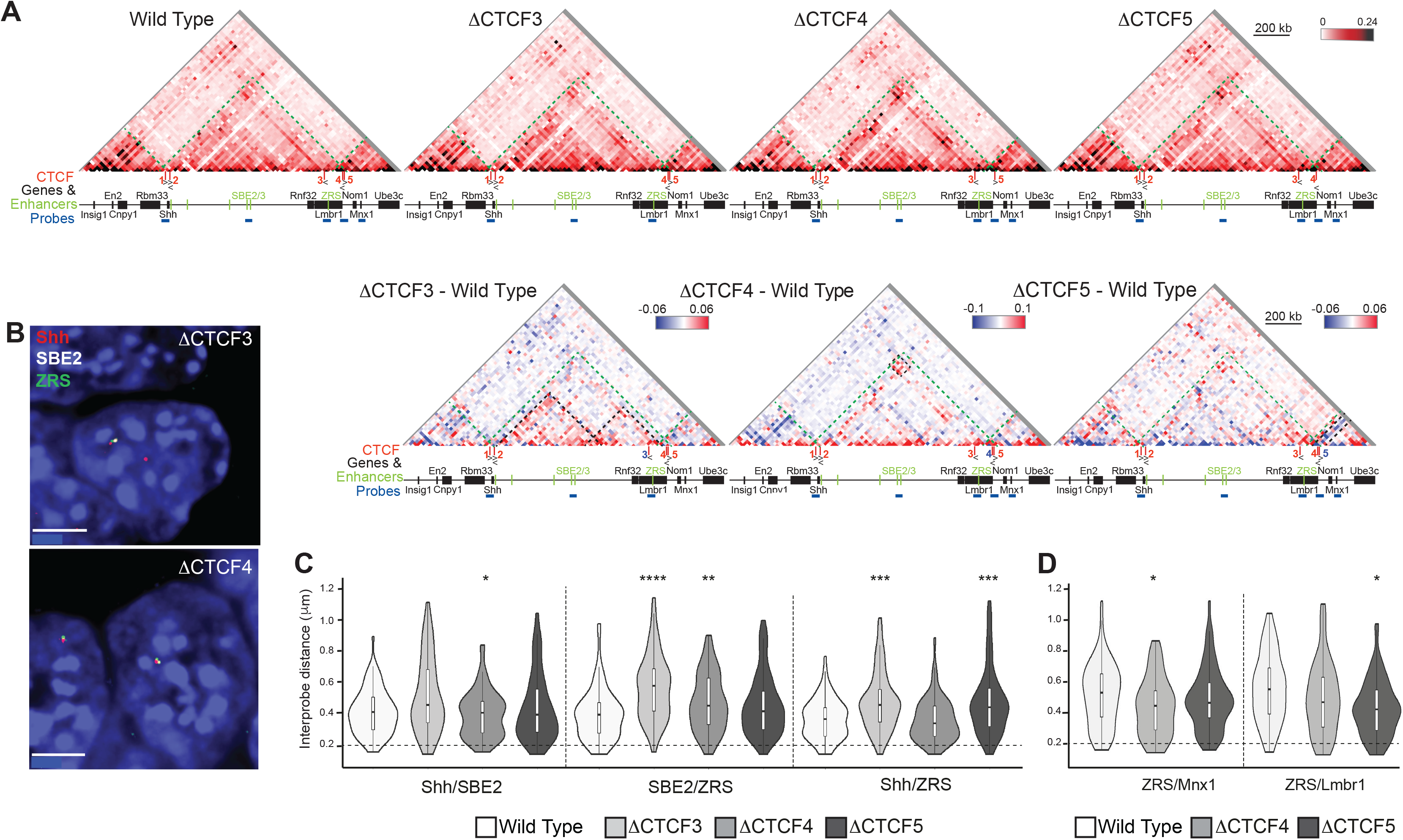
5C and 3D FISH identifies perturbations to local chromatin conformation in ΔCTCF3, ΔCTCF4 and ΔCTCF5 ESCs. **(A)** 5C heat maps from wild type, ΔCTCF3, ΔCTCF4 and ΔCTCF5 ESCs (top), across the 1.7-Mb *Shh* region shown in Figure 1A. Heat map intensities represent the average of interaction frequency for each window, colour-coded according to the scale shown. Interaction frequencies were normalized based on the total number of sequence reads in the 5C data set and the data shown is binned over 22.5-kb windows. Below are heat maps comparing ΔCTCF3, ΔCTCF4 or ΔCTCF5 enrichment (red) with wild type (blue). Green dashed lines indicate TAD boundaries, black dashed lines highlight enriched contacts within sub-TADs (ΔCTCF3), enriched contacts between genomic regions containing *Shh*/CTCF2 and ZRS/CTCF3 (ΔCTCF4) and loss of contacts between *Lmbr1* promoter/CTCF5 and the rest of the TAD containing *Mnx1* (ΔCTCF5). Data for biological replicates are in Supplemental Figure S4. **(B)** Images of representative nuclei from ΔCTCF3 and ΔCTCF4 ESCs showing FISH signals for *Shh*/SBE2/ZRS probes. Scale bars = 5 μm. **(C)** Violin plots show the distribution of interprobe distances (μm) between *Shh*/SBE2, SBE2/ZRS and *Shh*/ZRS probes in wild type, ΔCTCF3, ΔCTCF4 and ΔCTCF5 ESCs. Horizontal dashed line shows the proportion of alleles that are colocalised (< 200 nm). The statistical significance between data sets was examined by Mann-Whitney U Tests, * < 0.05, ** < 0.01, *** < 0.001, **** < 0.0001. **(D)** As in **(C)** but for ZRS/*Mnx1* and ZRS/*Lmbr1* probes in wild type, ΔCTCF4 and ΔCTCF5 ESCs.

Finally, we generated ESC lines with deletions of CTCF binding sites at the *Lmbr1* promoter (ΔCTCF4) and 5′ *Lmbr1* (ΔCTCF5), both of which are located at the boundary between the *Shh* TAD and the adjacent TAD containing *Mnx1* (Fig. 1A). Neither of these deletions affected *Shh* TAD integrity or boundary positions (Figures 4A & S4A, top row, centre-right and right-hand heatmaps). However, the CTCF3/ZRS genomic region had enriched interactions (red) across the *Shh* TAD in ΔCTCF4 ESCs, especially with the *Shh* locus itself (Figs. 4A & S4A, bottom row, middle heatmaps). The CTCF motif within CTCF5 is oriented towards the adjacent *Mnx1*-containing TAD and deletion of the site caused a loss of interactions (blue) between this boundary region and the *Mnx1* TAD (Figs. 4A & S4A, bottom row, right-hand heatmaps).

FISH revealed significantly increased inter-probe distances between *Shh* and ZRS in ΔCTCF5 cells and between SBE2 and ZRS in ΔCTCF4 cells, which contrasts with the 5C data (Fig. 4C). There are also decreased distances seen between ZRS and *Mnx1* in the adjacent TAD in the absence of CTCF4, something not apparent in the 5C data (Fig 4D). Also, in contrast to 5C data, loss of CTCF5 decreases distances between ZRS and *Lmbr1* promoter compared to wild type (Fig. 3D).

We conclude that deletion of CTCF binding sites at either of the *Shh* TAD and sub-TAD boundaries, especially CTCF1, 2 and 3, affects local chromatin organisation in ESCs and disrupted *Shh*/ZRS spatial proximity.

### Reduced *Shh*-ZRS colocalisation upon the loss of CTCF1, 2 and 3 in the limb

To test how disrupted TAD organisation impacts on chromosome conformation and *Shh* gene expression during embryonic development, we generated mouse lines carrying each of the CTCF deletions. We previously reported a significantly enhanced *Shh*-ZRS colocalisation in the Zone of Polarising Activity (ZPA) of the limb bud where *Shh* is active compared to non-expressing limb tissues. This colocalisation depends on a fully functional ZRS and, when mutations are made within ZRS that deleteriously affect *Shh* expression, colocalisation rates are dramatically reduced (Lettice et al., 2014; Williamson et al., 2016). Therefore, we assayed the spatial proximity of *Shh*, SBE2 and ZRS by FISH in E11.5 mouse embryo sections that include posterior (ZPA) and anterior distal limb tissue from wild-type and homozygous ΔCTCF mutant embryos (Fig. 5A).

**Figure 5.**
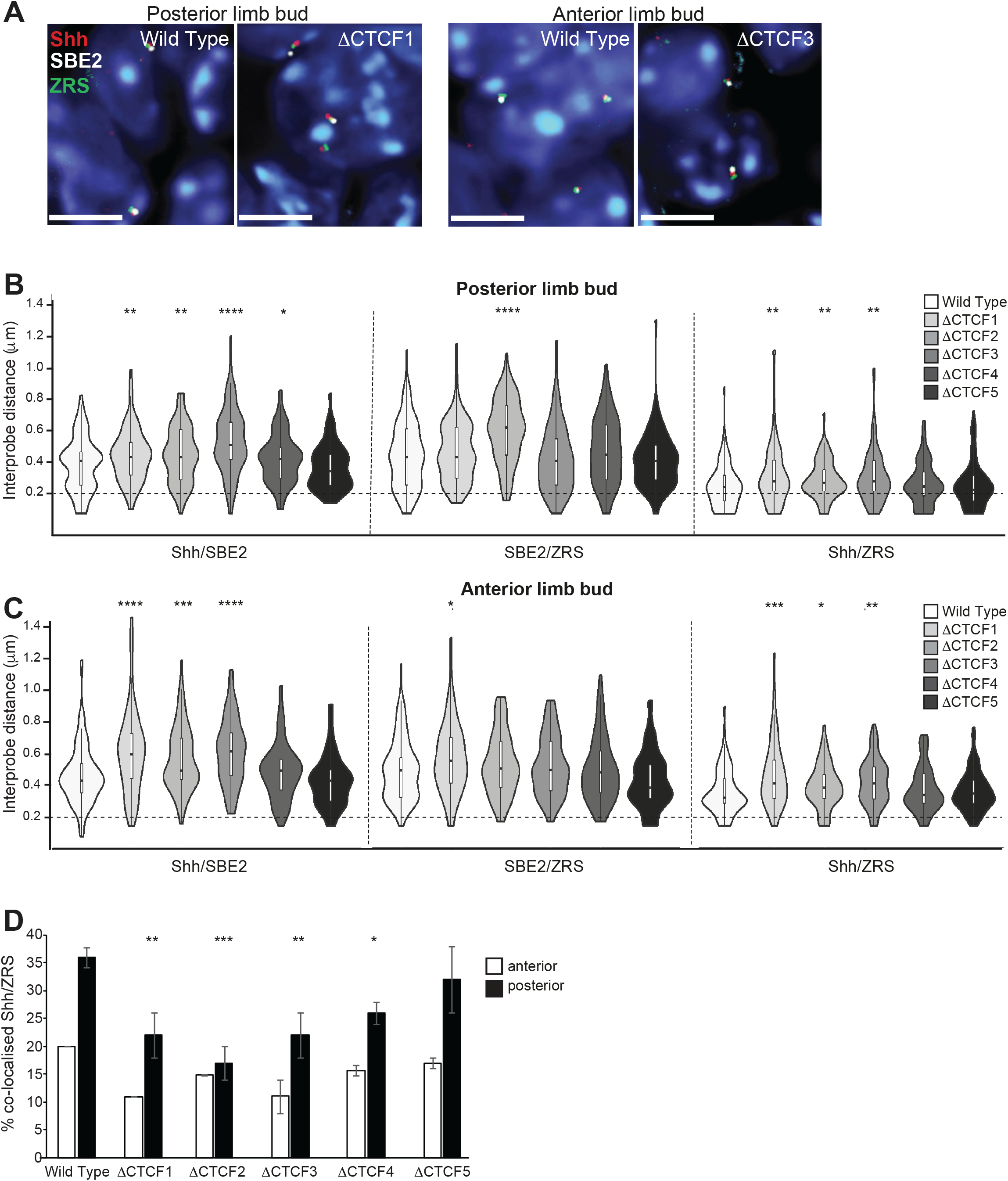
Perturbation of chromatin conformation within the *Shh* TAD in the distal limb bud of ΔCTCF mutant embryos. **(A)** Images of representative nuclei from E11.5 ZPA and distal anterior limb bud in wild type, ΔCTCF1 and ΔCTCF3 embryos showing FISH signals for *Shh*/SBE2/ZRS probes. Scale bars = 5 μm. **(B) & (C)** Violin plots show the distribution of interprobe distances (μm) between *Shh*/SBE2, SBE2/ZRS and *Shh*/ZRS probes in E11.5 wild type, ΔCTCF1, ΔCTCF2, ΔCTCF3, ΔCTCF4 and ΔCTCF5 in **(B)** ZPA and **(C)** distal anterior limb bud. Horizontal dashed lines show the proportion of alleles that are colocalised (< 200 nm). The statistical significance between data sets was examined by Mann-Whitney U Tests, * < 0.05, ** < 0.01, *** < 0.001, **** < 0.0001. **(D)** Histograms show the percentage of colocalised *Shh*/ZRS probe pairs (<200nm) in wild type and each of the ΔCTCF E11.5 embryos for distal anterior and ZPA limb bud tissue. Error bars represent SEM obtained from two or three different tissue sections. The statistical significance between data sets was examined by Fisher’s Exact Tests, * < 0.05, ** < 0.01, *** < 0.001.

In both regions of the wild type limb bud analysed (ZPA and anterior), *Shh*-ZRS distances were shorter, than between *Shh*-SBE2 and SBE2-ZRS, consistent with these two loci being maintained in spatial proximity across the limb bud (Figs. 5B & C, white violin plots). Distances between *Shh* and both SBE2 and ZRS were significantly increased in ΔCTCF1, 2 and 3, but not ΔCTCF4 and 5 embryos, similar to that observed in ESCs (Figs. 5B & C). This disruption of *Shh*-ZRS spatial proximity in the ZPA of ΔCTCF1, 2, and 3 mutant embryos significantly reduced the frequency of *Shh*-ZRS colocalisation (<200nm) down to levels seen in non-expressing parts of the wild-type limb bud (Fig. 5D).

### *Shh* expression patterns and development are unaffected in CTCF site deletion mice

Our data indicate that deletion of individual CTCF sites affects TAD boundaries, intra- and inter-TAD interactions and enhancer-promoter co-localisation frequencies. These alterations in 3D chromosome conformation would be predicted to have an effect on gene expression. Surprisingly, however, we found that mice homozygous for any of the ΔCTCF deletions are viable, fertile and have no detectable phenotype. In situ hybridisation in homozygous mutant embryos showed a normal pattern of *Shh* expression, and at similar levels to wild type, (Fig. 6A and B). At E11.5 expression is detected only within the developing midline of the brain, the Zli and the medial ganglionic eminence in the head and staining is visible in the floor plate and notochord, the ZPA of the limb buds and umbilicus in the body. No ectopic expression is detected at the midbrain / hindbrain junction driven by neighbouring *En2* or *Cnpy1* enhancers (Fig 6A), Conversely, in embryos homozygous for ΔCTCF1 and ΔCTCF2 there is no evidence for ectopic *En2* and *Cnpy1* expression in any of the normal sites of *Shh* expression in the brain (Fig 6C and D).

**Figure 6.**
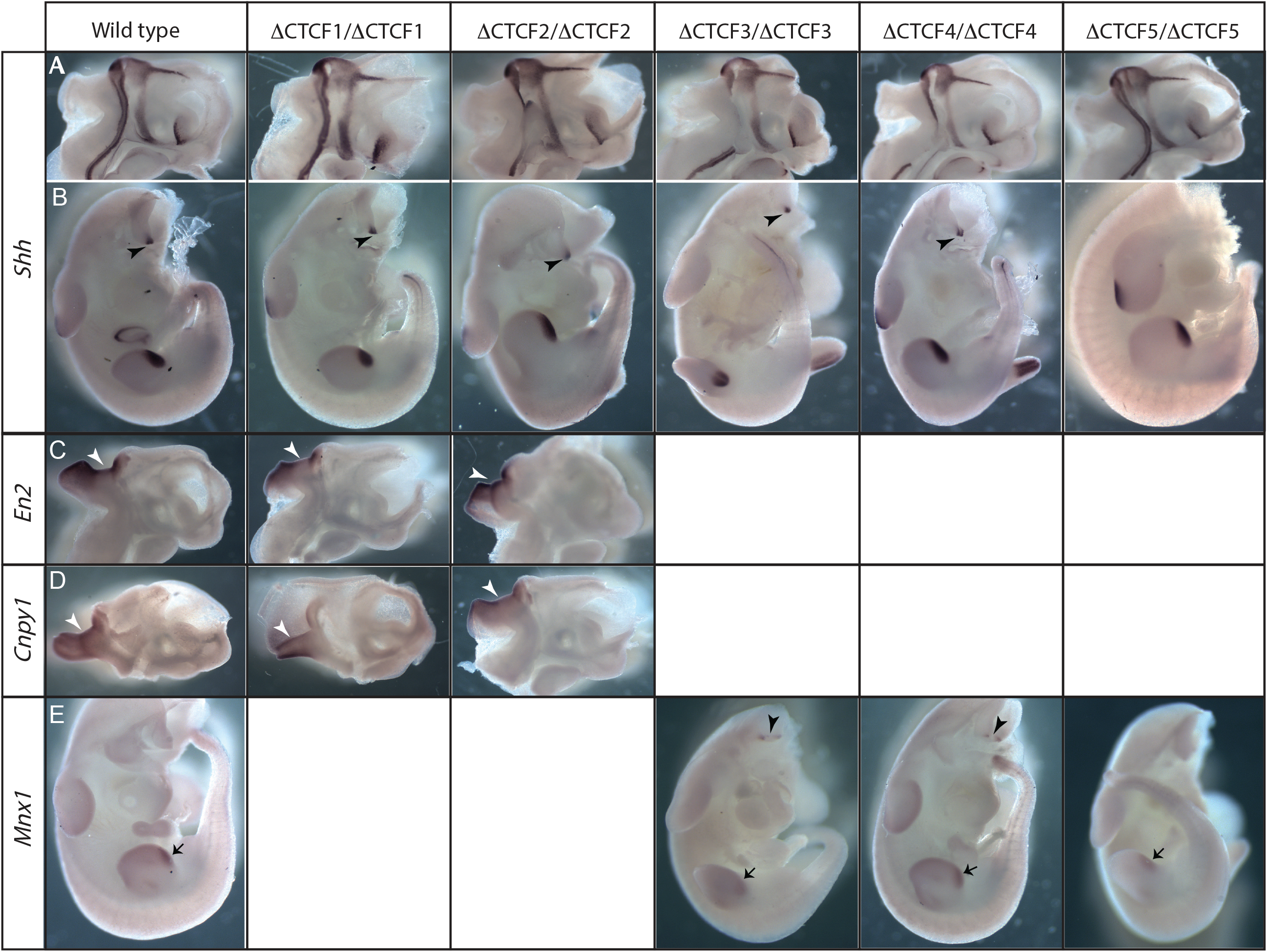
Expression patterns of *Shh* and genes in neighbouring TADs are unaffected by CTCF site deletions. In situ analysis of gene expression at E11.5 in wild type and embryos homozygous for each of the ΔCTCF lines. **(A and B)** show normal expression of *Shh* in the midline of bisected heads **(A)** and in the bodies **(B)**, expression is detected in the ZPA of the limb bud and in the floorplate and notochord (arrowheads). No ectopic expression is observed. **(C and D)** show expression of *En2* (**C**) or *Cnpy1* (**D**) in bisected heads. Expression is detected only at the mid brain hindbrain junction (arrowheads). **(E)** Expression of *Mnx1*in the ZPAs of the limb buds (arrows) and in the motor neurons (arrowsheads).

Similarly, no ectopic *Shh* expression is detected in motor neurons driven by *Mnx1* enhancers in the TAD beyond ZRS/*Lmbr1* (Fig. 6A), and *Mnx1* was not expressed ectopically in any of the normal sites of *Shh* expression in embryos carrying homozygous deletions of CTCF3, CTCF4 and CTCF5 (Fig 6E). These findings indicate that enhancer/promoter specificity is maintained in these deletion embryos and that there is no cross-talk across TAD boundaries resulting in ectopic expression driven by *Shh* enhancers, even in the absence of these CTCF sites.

Mice heterozygous for a *Shh* null allele express only about 60% wild type levels of *Shh* but develop normally, suggesting that there is a wide range of *Shh* expression levels capable of driving normal development. Indeed, in the limb *Shh* levels must fall to about 20% of wildtype before development becomes perturbed and digits are lost (Lettice et al., 2017). We therefore made compound heterozygotes carrying both the ΔCTCF1 and *Shh* null alleles to uncover subtle effects on *Shh* expression caused by the CTCF1 deletion. These *Shh*^ΔCTCF1/-^ mice develop normally and are viable and fertile, further suggesting that deletion of CTCF1 results in no deleterious changes in *Shh* expression.

### A 35kb deletion that removes the *Lmbr1* promoter and TAD boundary disrupts chromatin conformation with no deleterious phenotype

Deletion of CTCF1 3′ of *Shh* showed that this position was important for the TAD boundary location and ensuring *Shh* remained in physical proximity with its regulatory domain (Figs. 3 & S3), even though the loss of this site had no apparent phenotypic consequence (Fig. 6). Deleting either CTCF4 or CTCF5 at the other TAD boundary had no effect on the *Shh* boundary location and resulted in a minimal loss of proximity between ZRS and *Shh* (Figs. 4 & S4). Loss of TAD boundary regions can result in the merging of adjacent TADs and the ectopic activation of genes in one TAD by enhancers in the other merged TAD, with phenotypic consequences (Fabre et al., 2017; Lupiáñez et al., 2015). However, this involved the deletion of sizeable stretches of DNA across the boundaries in question, tens of kilobases rather than individual CTCF sites. In addition to CTCF binding sites, a number of features are found enriched at TAD boundaries including those associated with active promoters (Dixon et al., 2012). To determine if a more extensive deletion across the *Lmbr1* boundary results in the merging of adjacent TADs, a homozygous 35kb deletion (Δ35) was generated which removed CTCF4 and CTCF5, in all covering about 13kb upstream of the *Lmbr1* TSS, as well as TSS itself and first two exons of *Lmbr1* (Fig. 1A). RT-PCR confirmed that this deletion eliminates transcription throughout the 5′ end of *Lmbr1* in both isolated limb buds and the rest of the body (Fig 7A).

**Figure 7.**
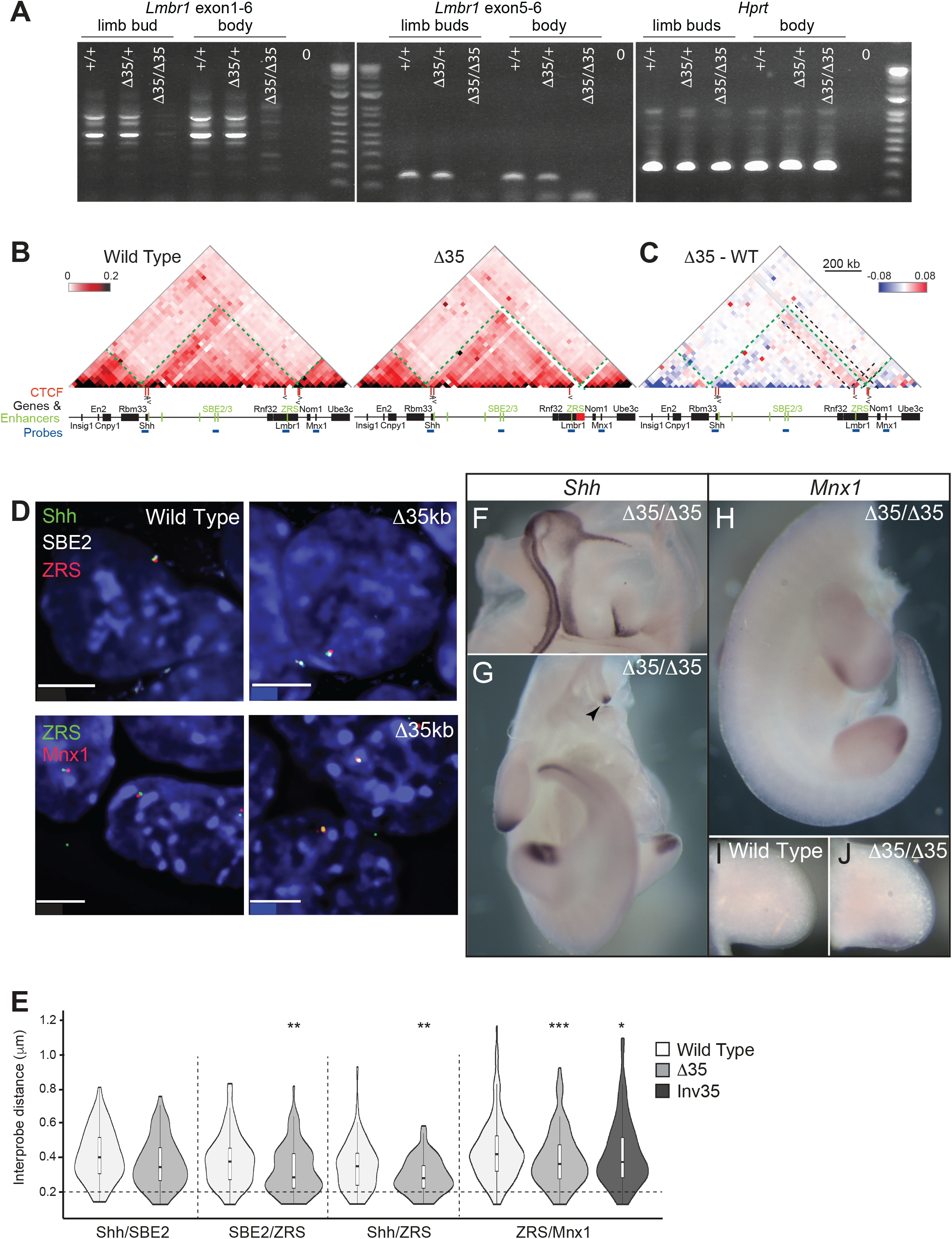
Chromosome conformation and gene expression as a consequence of a 35-kb deletion at the *Lmbr1* TAD boundary. **(A)** RT-PCR analysis of gene expression in limb buds and bodies from E11.5 wild type, heterozygous and homozygous 35kb deletion (Δ35) embryos showing a loss of transcription through *Lmbr1*. **(B)** 5C heat maps from wild type and Δ35-kb ESCs, across the 1.7-Mb *Shh* region shown in Figure 1A. Heat map intensities represent the average of interaction frequency for each window, colour-coded according to the scale shown. Interaction frequencies were normalized based on the total number of sequence reads in the 5C data set and the data shown is binned over 35-kb windows. **(C)** Heat map comparing Δ35-kb enrichment (red) with wild type (blue). Green dashed lines indicate TAD boundaries, black dashed lines highlight change in boundary position and enriched contacts between the genomic region containing ZRS/CTCF3 and the rest of the *Shh* TAD and across the perturbed TAD boundary up to *Mnx1* in Δ-35kb cells. **(D)** Images of representative nuclei from wild type and Δ-35kb ESCs showing FISH signals for *Shh*/SBE2/ZRS and ZRS/*Mnx1* probes. Scale bars = 5 μm. **(E)** Violin plots show the distribution of interprobe distances (μm) between *Shh*/SBE2, SBE2/ZRS, *Shh*/ZRS and ZRS/*Mnx1* probes in wild type and Δ-35kb ESCs, and ZRS/*Mnx1* distances in 35kb inversion ESCs. Horizontal dashed line shows the proportion of alleles that are colocalised (< 200 nm). The statistical significance between data sets was examined by Mann-Whitney U Tests, * < 0.05, ** < 0.01, ** < 0.001. **(F & G)** In situ hybridisations showing normal *Shh* expression in a bisected head and body, respectively, of a Δ35/Δ35 E11.5 embryo. **(H-J)** In situ hybridisations for *Mnx1* in a Δ35/Δ35 homozygote **(H)** and limb bud **(J)** and for comparison lower levels of staining in a wildtype limb bud is shown in **(I).** Staining in **(I)** and **(J)** was stopped before wildtype signal was apparent to highlight *Mnx1* up-regulation.

5C showed that Δ35 caused the relocation of the TAD boundary a further ∼40kb 5′ of the *Lmbr1* promoter towards the promoter of *Nom1* (Figs. 7B & S5A & B), rather than a merging of the adjacent TADs. There are enriched interactions of the region from CTCF3 to *Nom1* across the *Shh* TAD in Δ35 cells (red in Fig. 7C) compared to wild type and the CTCF3/ZRS genomic region gains interactions into the adjacent TAD up to *Mnx1* (Figs. 7C & S5B), whereas the region around *Nom1* had reduced interactions with its own TAD. Virtual 4C plots derived from the 5C data show that ZRS gains contacts with the rest of the *Shh* TAD and into the adjacent TAD up to *Mnx1* (Fig. S5C, top graphs), and the new boundary region gains contacts with the *Shh* TAD while losing interactions with the *Mnx1* TAD (Fig. S5C, bottom graphs). 3D-FISH (Fig. 7D) showed that distances between ZRS and the other three labelled loci (*Shh*, SBE2 and *Mnx1*) were all significantly decreased in Δ35 (Fig. 7E), which corresponds to the 5C and virtual 4C data. The reduced spatial distance between ZRS and *Mnx1* was not due to the linear genomic distance being reduced because of the 35kb deletion, as similar effects were seen in cells carrying an inversion of this DNA rather than a deletion (Fig. 7E).

Removal of the *Shh* TAD boundary at the *Lmbr1* promoter relocates the boundary to the promoter of *Nom1*, and the ZRS has enhanced ability to contact sequences both within its own TAD and the *Mnx1* TAD. Despite these differences, Δ35 homozygous mice were viable, fertile and had no apparent phenotype. The *Shh* expression pattern is also indistinguishable from wild type (Figs. 7F & G) - in particular midline expression is detected in the floor plate and notochord as one stripe down the body (Fig 7G, arrow head), with no evidence for expression as two more lateral stripes driven by *Mnx1* motor neuron enhancers (Fig. 6E). Even in the sensitised background of the *Shh* null chromosome compound *Shh*^Δ35/-^ mice are also phenotypically normal.

Interestingly however, given the decreased distances measured by FISH between ZRS and *Mnx1*, the limb expression of *Mnx1* was increased in *Shh*^Δ35/Δ35^ embryos in comparison to wild-type embryos (Figs. 7H-J). This suggests that this deletion, that encompasses the TAD boundary, enhances the ability of the *Mnx1* promoter to respond to the ZRS. However, no upregulation of *Mnx1* expression is seen in the pharyngeal endoderm and developing lungs which would be driven by the enhancers neighbouring ZRS, MACS1 and MFCS4.

## DISCUSSION

A systematic genetic approach to remove most of the *Shh* enhancers (excluding ZRS) (Δ700), to delete individual CTCF sites, and to delete or invert large regions encompassing a TAD boundary, has enabled us to use chromosome conformation capture and imaging to assay the resulting perturbations to chromosome organisation within the *Shh* regulatory TAD, and between this and neighbouring TADs. Despite this, we detected little or no perturbation of gene regulation during embryonic development and no detectable phenotype in animals that can be attributed to this altered chromosome conformation.

### ZRS activity is not distance dependent and does not require factors located within the intervening gene desert

5C analysis confirmed that TAD boundaries were unaffected by removal of most of the internal region of the *Shh* TAD (Δ700) (Figs. 1 & S1), with *Shh* and its remaining enhancers still located within the same, but smaller, TAD. This large deletion did cause extensive disruption to the developing embryo, mainly, it can be assumed, due to the loss of several known forebrain and epithelial enhancers within the deleted region. However, even in embryos homozygous for the 700kb deletion, which relocates ZRS to less than 100kb distant from *Shh*, ZRS function is maintained, there is no detrimental effects on limb bud-specific *Shh* activation and normal development of the limbs occurs. Therefore, the large genomic distance from *Shh* is not intrinsic to the function of the ZRS. This is in contrast to the loss of interactions following similar perturbations between a limb-specific enhancer and *Hoxd13* that resulted in loss of *Hoxd13* activity (Fabre et al., 2017).

### Loss of CTCF sites at the *Shh* TAD boundaries disrupts chromatin architecture, and impacts *Shh*/ZRS spatial proximity

We have previously shown that *Shh* and ZRS are in spatial proximity (∼300nm) in the early embryo in both expressing limb tissue and the non-expressing adjacent flank (Williamson et al., 2016). Here, using 5C on cells dissected from E11.5 anterior and posterior limb buds we show that this is driven by a looping interaction between the sites 3′ and 5′ of *Shh* (containing CTCF1 and CTCF2 sites) and a region within intron 5 of *Lmbr1* about 20kb from ZRS (CTCF3) (Figs. 2 & S2). This loop is also present in ESCs, and spatial proximity of *Shh* and ZRS is lost upon the deletion of any one of the three CTCF sites in both ESCs and E11.5 limb bud tissue (Figs. 3, 4, 5, S3 & S4). Deleting CTCF sites at the *Lmbr1* promoter TAD boundary (CTCF4 and CTCF5) had less effect on *Shh*/ZRS spatial proximity. Increased inter-probe distances between either *Shh* or ZRS and the forebrain enhancer SBE2 located at the centre of the TAD suggest that the loss of spatial proximity may be due to a general decompaction throughout the TAD.

### *Shh* responds to its developmental enhancers regardless of TAD disruption

5C analysis in ESCs suggests that the disruption caused by the deletions removes *Shh* from its regulatory TAD (ΔCTCF1) or re-enforces contacts within sub-TAD domains such that the forebrain enhancers are sequestered in one and ZRS and the long-range epithelial enhancers in the other, with a loss of interactions between both sub-TADs and either sub-TAD with *Shh* (ΔCTCF2 and ΔCTCF3). Nevertheless, in all of these configurations, the pattern of *Shh* during embryonic development appears to be normal and the resulting mice have no detectable phenotype. This indicates that communication between *Shh* and its extensive set of developmental enhancers is remarkably robust to TAD perturbation.

### Ectopic expression across disrupted TAD boundaries is not common

Loss of CTCF1 not only moves the TAD boundary ∼40kb to beyond the 5′ end of *Shh* but also enables greater interactions between *Shh* the adjacent TAD which contains other genes and their enhancers active during brain development, but in a pattern distinct from *Shh*. *En2* is expressed at the mid-hindbrain boundary, a pattern at least partly dependent on an enhancer binding Pax2/5/8 (Li Song and Joyner, 2000). Similarly, *Cnpy1* expression at the mid-hindbrain boundary is thought to be important for FGF signalling (Hirate and Okamoto, 2006). Despite increased chromatin interactions over the *Shh* TAD boundary in ΔCTCF1, there is no ectopic expression of *Shh* in the mid-hindbrain driven by the *En2*/*Cnpy1* enhancers and, vice versa, there is no ectopic expression of *En2*/*Cnpy1* at sites driven by *Shh* enhancers (Fig. 6).

The *Lmbr1* boundary has been suggested to be less precise than the *Shh* boundary from a structural and regulatory point of view (Symmons et al., 2016). Indeed, deletion of CTCF4 or 5 had little effect on this *Shh* TAD boundary. However, ΔCTCF5 weakened the boundary of the neighbouring *Mnx1* TAD and increased proximity between ZRS and *Mnx1* was detected in ΔCTCF4. However, in neither case was there evidence for enhanced expression of *Mnx1* – e.g. in limb buds driven by ZRS – beyond that detected in wild-type embryos. Interestingly, even in wildtype situations, *Mnx1* has a weak expression domain concomitant with the limb bud ZPA, suggesting that this gene may be influenced by ZRS activity emanating from the adjacent TAD. Nor was there evidence of the *Mnx1* motor-neuron enhancer (Zelenchuk and Brusés, 2011) driving expression of *Shh* in motor neurons of the developing neural tube in any of the mutant embryos.

A larger (35kb) deletion of this boundary removing CTCF4, CTCF5 and the promoter/first two exons of *Lmbr1*, enhanced ZRS 5C contacts across both the *Shh* TAD and into the neighbouring *Mnx1* TAD (Figs. 7 & S5). Increased *Mnx1* expression in the ZPA of embryos homozygous for the 35kb deletion suggests that the potentially increased contacts between *Mnx1* and ZRS identified in ESCs could be enabling greater activation of this gene by the *Shh* limb enhancer.

### Perturbations of the *Shh* TAD boundaries can negatively impact on gene-enhancer co-localisation but are insufficient to cause a deleterious phenotype

It is commonly thought that enhancer driven gene-activation required ‘contact’ or very close apposition of the enhancer and promoter. Inversions encompassing the *Shh* TAD boundaries that disrupted TAD integrity and significantly increased the genomic distance between *Shh* and ZRS result in severe limb malformations, suggesting that these rearrangements prevent ZRS from contacting/regulating the *Shh* promoter (Symmons et al., 2016). These data and our 5C and FISH analysis which shows that the *Shh* TAD forms a compact, discrete regulatory hub (Williamson et al., 2016) suggest that 3D organisation of the *Shh* TAD could allow distal enhancers to come into close proximity to selectively regulate *Shh* expression. However, in the functionally relevant cells of the limb bud ZPA, ZRS colocalisation (<200 nm) with *Shh* was reduced to levels of the non-expressing distal anterior levels in ΔCTCF1, ΔCTCF2 and ΔCTCF3 homozygous embryos without adversely affecting *Shh* expression (Fig. 6) and with no subsequent phenotypical effects. We have previously shown that reduced *Shh*-ZRS colocalisation as a consequence of deleting the 3′ end of ZRS (which was shown to be not required for activating proximal expression of a reporter gene in the ZPA) caused the loss of *Shh* forelimb bud expression and severe attenuation of hindlimb bud expression which resulted in a range of limb malformations (Lettice et al., 2014). Therefore, loss of colocalisation on its own does not have severe enough effects on levels of expression to result in limb malformations.

All embryos homozygous for one of the five CTCF binding domain deletions and even with the 35kb deletion of the *Lmbr1* boundary developed normally and were able to reproduce. Moreover, sufficient expression of *Shh* was maintained in compound heterozygote embryos carrying either ΔCTCF1 or the 35kb deletion opposite a *Shh* null allele for these mice to have no phenotype. A contemporaneous study on the same genomic territory has largely re-capitulated these results – deletions of *Lmbr1* CTCF sites and the gene promoter caused perturbations to local chromatin conformation but *Shh* expression, although reduced, was enough to drive normal limb development (Paliou et al., 2019). The *Shh* regulatory landscape is set up to ensure optimal activation of the gene and here we have shown this is robust to perturbations of TAD integrity and structure. Only large-scale disruptions incorporating boundaries appear to cause TADs to merge with resulting developmental defects (Lupiáñez et al., 2015).

Our data suggest that TADs are largely structural and play no overt role in regulating gene expression. We speculate that the largely unvarying organisation of TADs could have provided the necessary stable genomic environment for the accumulation of regulatory elements over evolutionary time rather than being essential for target gene activation.

## Acknowledgements

We thank the staff of the IGMM advanced imaging resource and technical services for their assistance with imaging and sequencing. We would also like to thank the staff at the BRF/Evans Building for expert technical assistance. This work was supported by the Medical Research Council, UK.

